# Fast slow folding of an Outer Membrane Porin

**DOI:** 10.1101/2021.04.06.438691

**Authors:** Eve E. Weatherill, Monifa A. Fahie, David P. Marshall, Rachel A. Andvig, Matthew R. Cheetham, Min Chen, Mark I. Wallace

## Abstract

In comparison to globular proteins, the spontaneous folding and insertion of *β*-barrel membrane proteins is surprisingly slow, typically occurring on the order of minutes. Using single-molecule Förster Resonance Energy Transfer to report on the folding of fluorescently-labelled Outer Membrane Protein G we measured the real-time insertion of a *β*-barrel membrane protein from an unfolded state. Folding events were rare, and fast (<20 ms); occurring immediately upon arrival at the membrane. This combination of infrequent, but rare, folding resolves this apparent dichotomy between slow ensemble kinetics, and the typical timescales of biomolecular folding.

## 1. Introduction

It is well established that water-soluble proteins rapidly fold through a funnel-like energy landscape to the lowest energy state via a collection of pathways [1, 2, 3, 4]; but in comparison, the folding mechanisms of membrane proteins [5, 6, 7], and in particular *β*-barrel proteins [8, 9], are relatively poorly understood. *β*-barrel proteins are specific to the outer membranes of chloroplasts, mitochondria and gram-negative bacteria, playing a crucial role in processes as varied as energy production [10], photosynthesis, nutrient transport [11], enzymatic activity (e.g. protease [12] and lipase [13]), cellular adhesion [14], membrane anchoring [15], complement binding [16] and drug efflux [17]. Arguably the most important *β*-barrel proteins are the Bacterial Outer Membrane Proteins (OMPs) that, as their name suggests, reside in the outer membrane of gramnegative bacteria. Most OMPs form pores of some kind [8], and are characterised by a cylindrical topology, high thermodynamic stability, and (typically) an even number of *β*-strands. Highly conserved across gram negative bacteria, OMPs act as gatekeepers between the bacterium and its environment, and are thus promising candidate targets in the development of new antibiotics [18].

The biogenesis of OMPs *in vivo* is distinct from that of the *α*-helical transmembrane proteins of the inner membrane: Unfolded OMPs enter the periplasm via the Sec translocon, where they are protected from aggregation by a variety of molecular chaperones [19]. Folding and insertion into the outer membrane is assisted by the *β*-barrel assembly machinery complex (BAM) [20, 21, 22], which is thought to help OMPs overcome the energetic barrier to folding by local destabilisation of the outer membrane [23]. Although BAM acts to accelerate OMP folding kinetics in a manner analogous to classical chaperones, *β*-barrel folding and insertion also occurs spontaneously [24]. Much of our understanding of the folding of *β*-barrel proteins is built on in *vitro* studies in the absence of BAM, in which the folding of OMPs from denaturants into lipid membranes or detergent micelles can be measured precisely. The overall OMP folding pathway has been reviewed thoroughly elsewhere [9, 8]; but in summary, the spontaneous folding and insertion of *β*-barrel proteins follows a sequential pathway involving the rapid formation of a collapsed state before membrane absorption and then insertion of *β*-hairpins [25, 26]. The available biochemical and biophysical evidence supports a concerted mechanism, whereby the final folding step and membrane insertion occur simultaneously [27, 28, 29, 30]. There is, however, evidence that for some OMPs insertion occurs via intermediate states [31]. The membrane itself plays a key role in the kinetics of folding and insertion of OMPs: folding rates are dictated by membrane fluidity, thickness, curvature and head-group composition [24, 32, 28, 8, 33, 34, 35]. Overall, the picture for OMPs spontaneously folding and inserting into lipid membranes is one where kinetics are remarkably slow, typically in the order of minutes [32].

Although OMP folding kinetics are slow, it seemed implausible to us that the folding event itself could take so long at a molecular scale. To gain further insight we turned to single-molecule methods, which allow for the interrogation of heterogeneous molecular processes, and provide the means to resolve individual kinetic steps without the need to synchronise events.

Single-molecule Förster resonance energy transfer (smFRET) has proved to be an invaluable tool to measure protein folding kinetics over a wide range of timescales and protein systems [36, 23]. Confocal smFRET measurements have been particularly insightful; enabling the examination of folding dynamics of cytoplasmic [37, 38, 39, 40, 41] and membrane [42, 43, 44, 45] proteins. These methods have recently been applied to the folding and aggregation states of OMPs mediated by detergent [46, 47] and chaperonins [48]. However, due to the fast diffusion in solution, confocal single-molecule measurements generally lack the ability to directly observe transitions between such states. In contrast, continuous measurements of immobilised, confined, or encapsulated molecules permit transitions between conformational states to be measured [49, 50] (typically in exchange for time resolution). Continuous smFRET measurements have been used with great effect to probe functional dynamics [51, 52, 53, 54] and association dynamics [55, 56, 57, 58] of membrane proteins in lipid bilayers. However, their application to membrane protein folding in a lipid bilayer environment is, to our knowledge, yet to be demonstrated; such a study was recently proposed by Krainer et al [59] and here we seek to address that call.

To study single-molecule *β*-barrel folding we identified the *E. coli* outer membrane protein G (OmpG) [60] as a promising candidate. OmpG is a 14-stranded *β*-barrel notable for its monomeric status and flexible loops at the entrance to the pore [61, 62]. In comparison with trimeric OMPs, these characteristics simplify its folding landscape and have led to applications in nanopore sensing [61, 63, 62]. Detergent-mediated refolding of OmpG from a urea-unfolded state has been well characterised [64, 65], and kinetics of refolding in lipid vesicles, also in the presence of the BAM complex, occur on the order of minutes [66]. Using FRET-labelled OmpG we report on *β*-barrel folding into 1,2-dicetyl-*sn*-glycero-3-phosphocholine (DCPC) model membranes from urea using single-molecule Total Internal Reflection Fluorescence (smTIRF) microscopy.

## Methods

Detailed materials and methods are provided as Supplementary Information.

### Expression, purification and labelling of OmpG-2xCys

A double cysteine mutant of full-length OmpG (E2C and C281) was produced by site-directed mutagenesis and confirmed by DNA sequencing. Constructs were expressed and purified using a previously described method [69]. Samples were stored in denaturing buffer (8M Urea, 250 mM NaCl,1mM TCEP, 25 mM Tris.HCl (pH 8.0) at −80°C until required.

### Fluorescent labelling of OmpG Cy3/Cy5

OmpG-2xCys was purified by gel filtration in order to remove TCEP. Labelling was performed immediately afterwards by incubation with a 10-molar excess each of Cy3- and Cy5-maleimide for 60 min at room temperature. Excess label was removed by gel filtration. OmpG-Cy3Cy5 was stored at −80°C until required.

### Electrophysiology

For single-channel recordings, OmpG-Cy3Cy5 was refolded in detergent micelles and then reconstituted in a droplet interface bilayer (DIB). Droplet interface bilayers were produced as described in [**?**]. Voltage-clamped recordings of ionic current were made at room temperature and digitized at a rate of 1 kHz.

### Supported lipid bilayers

SLBs were prepared on glass coverslips by vesicle fusion [**?**] from SUVs consisting of 1.77 mM DCPC with 1.0mol% PEG(5K)-DPPE (and 3×10^−6^mol%TR-DHPE for lipid tracking experiments). For injection measurements, OmpG-Cy3Cy5 was diluted to 7 pM in denaturing buffer (8M urea, 250 mM NaCl, 10 mM tris pH 7) then 2 μl was added to the bulk solution during image acquisition. For SLBs containing pre-folded OmpG-Cy3Cy5, the protein stock was diluted to 700 pM in denaturing buffer, then 5 μl was mixed with 45 μl SUV stock and incubated for 1 hour at 37°C prior to SLB formation. For lipid tracking measurements, SLBs were imaged before and 10 minutes after the addition of OmpG Cy5, as described for OmpG-Cy3Cy5 injection.

## Results

### FRET-labelled OmpG reports on β-barrel folding

Maleimide-functionalised Cy3 (donor) and Cy5 (acceptor) fluorophores were stochastically conjugated to two engineered cysteine residues on the otherwise cysteine-free native OmpG. Labelling sites near the N- and C-termini, were selected in order to optimise the change in FRET efficiency between unfolded and folded states (Figure 1a).

**Figure 1.**
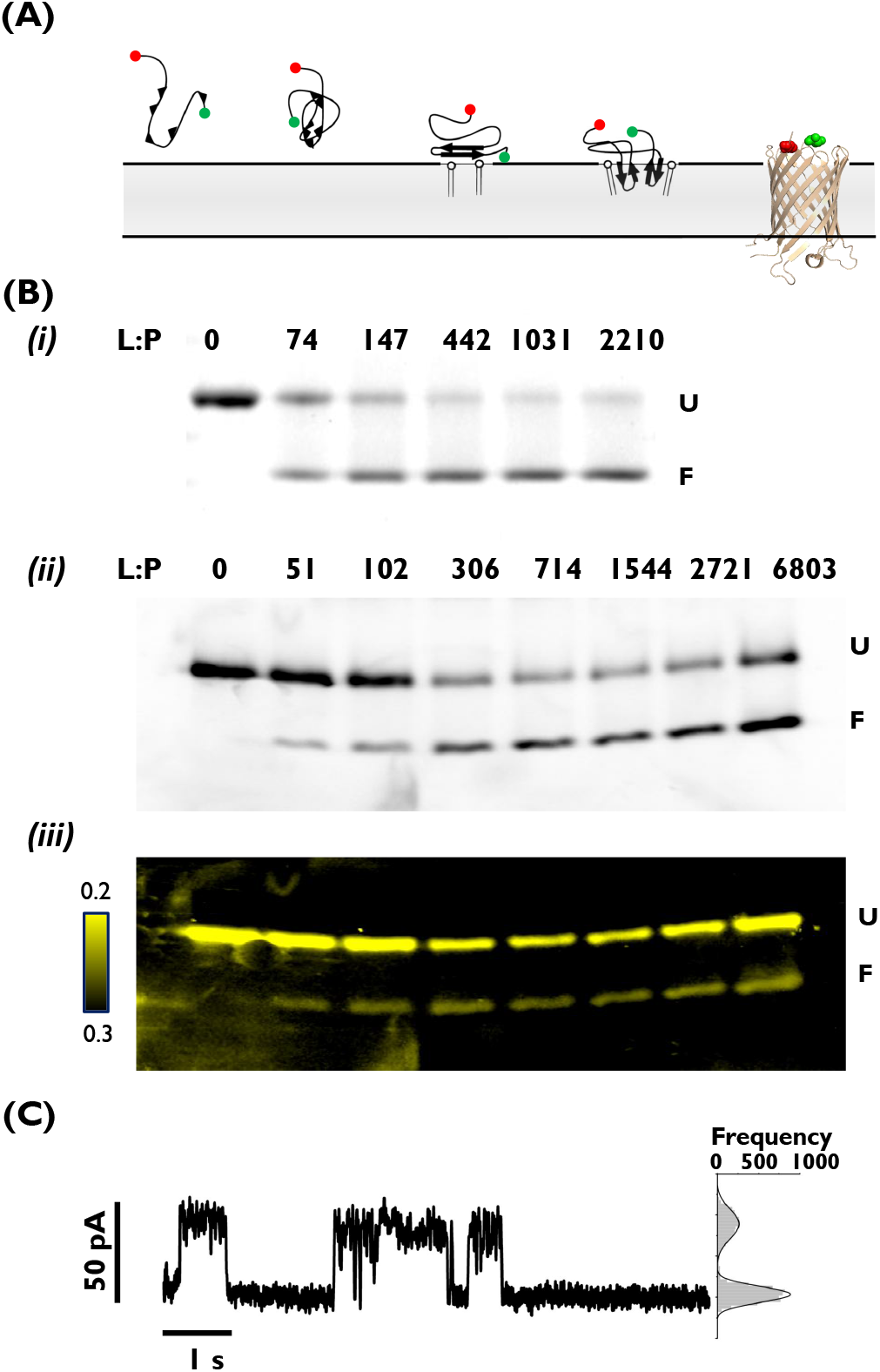
Doubly-labelled OmpG-Cy3Cy5 for probing *β*-barrel folding. (A) Model of *β*-barrel folding and membrane insertion as originally proposed in [25], modified to include crystal structure of OmpG (PBD:2F1C) with locations of fluorophore attachment near N- and C-termini highlighted red and green. (B) Urea-unfolded (i) OmpG-2xCys and (ii) OmpG-Cy3Cy5 after 1 h incubation with DCPC SUVs in 0.8 M urea, varying L:P ratio. The upper band (35 kDa) is the unfolded, membrane-associated state and the lower band (28 kDa) is the folded, native state. Gels (i&ii) were stained with coumassie, or (iii) illuminated within the excitation peak of the donor fluorophore and imaged with bandpass filters corresponding to donor (’D’) and acceptor (’A’) emission wavelengths. (Image shown is A/(D+A)). (C) Single-channel current recording of OmpG-Cy3Cy5, reconstituted in a DPhPC droplet interface bilayer.

Since OMPs are known to spontaneously fold at a higher rate and to a greater extent in bilayers with short hydrocarbon chains [32, 66], we refolded from 8M urea into small unilamellar vesicles (SUVs) of the short-chain (10-carbon) phospholipid DCPC. The extent of folding was assessed by both spectrometry (Figure 2b) and cold SDS-PAGE band-shift (Figure 1b): folded OMPs retain their structure in the presence of SDS at room temperature, and migrate faster than their unfolded counterparts because the folded state has a more compact structure [67]. Both (labelled) OmpG-Cy3Cy5 and (unlabelled) OmpG-2xCys displayed a lipid-dependent band-shift from the unfolded to folded state, characteristic of OMPs. OmpG-Cy3Cy5 FRET was also visualised directly in the SDS-PAGE gel where the efficiency of the (lower) folded protein bands is, as expected, consistently higher than the (upper) unfolded protein bands.

**Figure 2.**
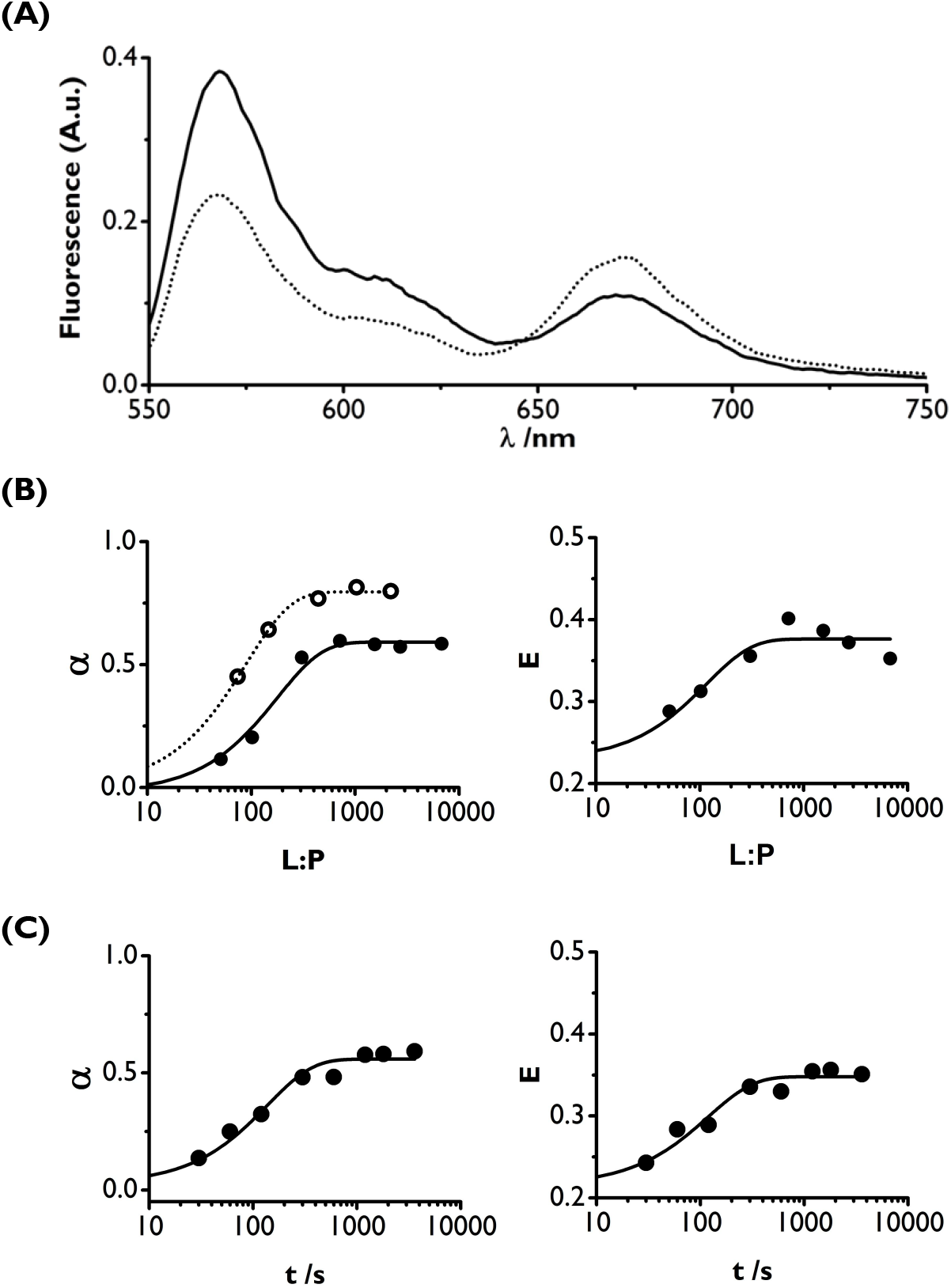
Titration and timecourse of OmpG-Cy3Cy5 folding into DCPC SUVs. Urea-unfolded OmpG after dilution to 0.8 M urea and incubation in the presence of DCPC SUVs. (A) Emission spectra of OmpG-Cy3Cy5 after 1 h incubation; L:P = 0 (solid line) and L:P = 714 (dotted line). (B) Folded fraction (left), and FRET efficiency (right) of OmpG-Cy3Cy5 (filled circles) and OmpG-2xCys (unfilled circles) after 1 hour incubation at varying L:P, with single exponential fit. (C) Folded fraction (left), and FRET efficiency (right) of timecourse of OmpG-Cy3Cy5 folding, with single exponential fit (*τ* = 82 ± 18 s and *τ* = 95 ± 18 s); L:P = 6250.

Electrophysiology was also performed to assess the functional state of OmpG following labelling. OmpG-Cy3Cy5 was refolded first in detergent and then reconstituted into a Droplet Interface Bilayer (DIB) following our previously published protocol [68]. We observed OmpG-Cy3Cy5 switching between open and closed states at neutral pH, with a conductance of 0.7 nS (Figure 1c), consistent with previous reports by both ourselves [69, 70] and others [65, 71, 72], indicative that the construct is capable of folding and forming a functional, native state.

Further ensemble measurements sought to establish whether FRET efficiency could be used as a direct readout for the folded state of OmpG-Cy3Cy5. An increase in FRET was observed when OmpG-Cy3Cy5 was diluted in the presence of SUVs compared to an SUV-free sample, denoted by a shift in donor and acceptor peaks in the emission spectra (Figure 2a). Titrating the L:P as before, and correcting the spectra for FRET-independent emission from both fluorophores (Figure S3), the FRET efficiency increased from 0.25 to 0.4 with a single exponential dependence on L:P ratio (*τ*_L:P_ = 177 ± 26) (Figure 2b). Quantification of the folded fraction of the same samples by densitometry analysis of the band-shift showed a similar dependence on L:P ratio (*τ*_L:P_ = 115 ±35). By comparison, (unlabelled) OmpG-2xCys appeared to fold more readily at lower L:P than OmpG-Cy3Cy5 (*τ* = 89 ± 5), and with a higher overall efficiency (*α*_max_ = 0.8 vs. 0.6).

A timecourse of OmpG-Cy3Cy5 folding in the presence of DCPC SUVs was performed, where folding was quenched periodically with SDS and assessed by band-shift assay and FRET efficiency (Figure 2c). The same range of folded fractions and FRET efficiencies were observed as with the L:P titration. Urea-unfolded OmpG-Cy3Cy5 transitioned to a folded, high-FRET state in the presence of DCPC SUVs with a halftime of approximately 90 seconds. This is consistent with previous studies of OMPs folding in the presence of short-chain lipids, which occurs on the order of minutes, and can follow multiple kinetic regimes [32, 66]. Thus, initial characterisation serves to confirm that OmpG-Cy3Cy5 forms functional channels, folds spontaneously into bilayers under equilibrium conditions at the expected rate, and that FRET efficiency is indeed a good readout to assess folded state.

### Folded OmpG gives rise to a single high-FRET state

Having established a FRET reporter of OmpG folding, we adapted our assay to observe folding events at the single-molecule level. Supported lipid bilayers (SLBs) were formed from DCPC SUVs and imaged by total internal reflection (TIRF) microscopy. 1 mol% PEG-DPPE was added to create a 4.5 nm hydrated cushion beneath the SLB [73, 74]; this distance exceeds the length of OmpG periplasmic loops [75].

To determine what FRET signals to expect from folded OmpG-Cy3Cy5 we first reconstituted OmpG-Cy3Cy5 directly into SUVs, before forming a SLB from the resulting proteoliposomes (Figure 3a). A L:P of 2.2 × 10^6^ gave a spot density suitable for single-molecule imaging. Fluorescence from donor and acceptor fluorophores was captured simultaneously on the same sensor. Mobile spots were visible in both channels, with a greater number in the donor channel (Figure 3c). Spots in the acceptor channel were only visible where FRET occurred, as the acceptor fluorophores were not excited directly. These correspond to doubly-labelled OmpG in a conformation in which the N- and C-termini are brought together. Acceptor-only labelled species (approx. 0.25 of the population) are not detected. Single molecule trajectories were selected for good signal:noise, donor and acceptor anticorrelation, single-step photobleaching, and non-crossing of other diffusing tracks. As expected, all selected trajectories exhibited a single high FRET state (*E* ≈ 0.910 ± 0.004, Figure 3D), before either donor (Figure 3bi) or acceptor photobleaching (Figure 3bii-vii). The lateral diffusion coefficient of these high-FRET spots *D*_lat_ = 0.072 ± 0.004 μm^2^ s^−1^, falls within the typical range for integral membrane proteins in polymer-cushioned SLBs [76, 77] (Figure 3E).

**Figure 3.**
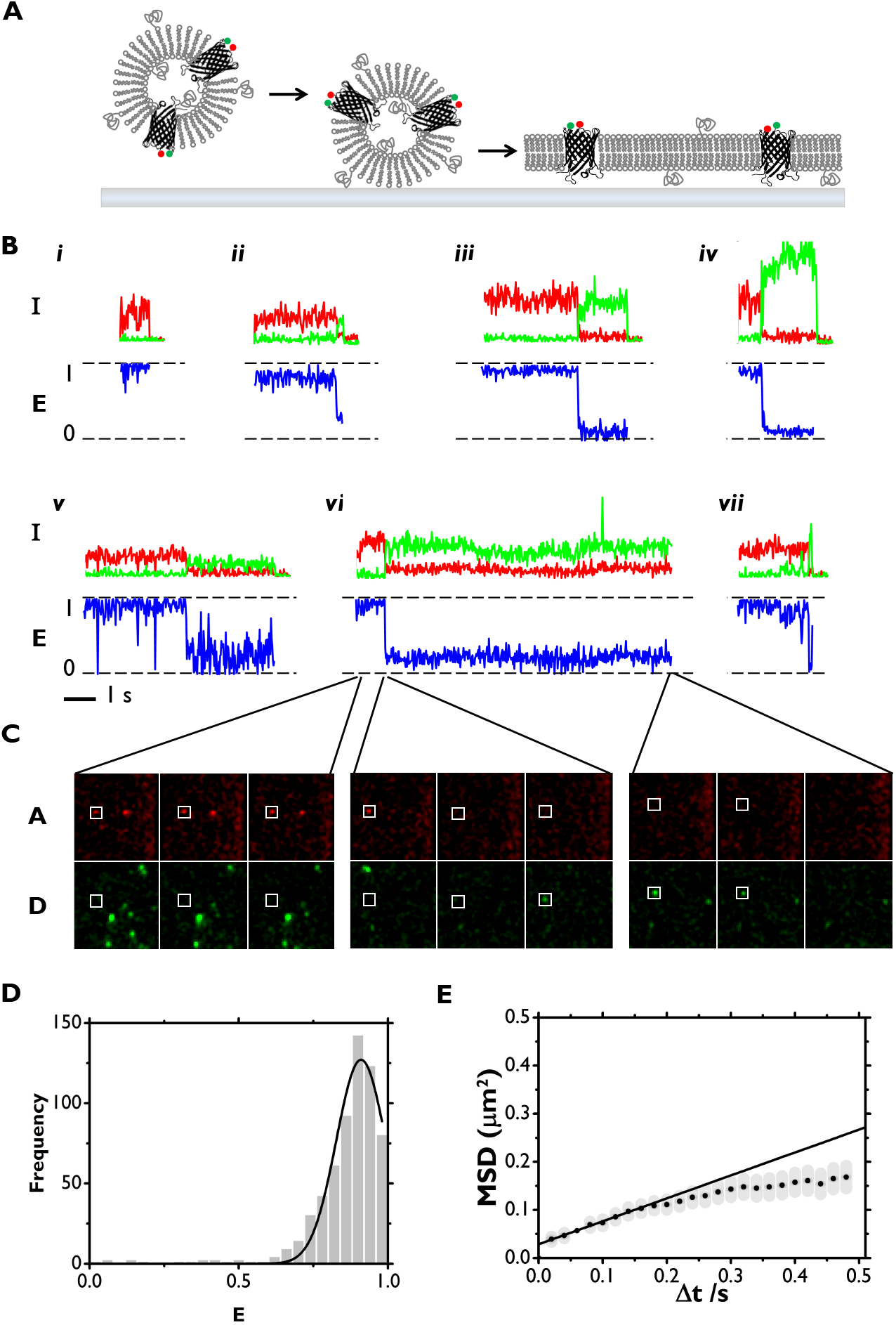
Pre-folded OmpG in DCPC SLBs: smFRET and SPT. (A) OmpG-Cy3Cy5 reconstituted into DCPC SUVs were fused onto glass, forming a supported lipid bilyer containing pre-folded OmpG. (B) Representative trajectories of tracked spots. Upper: Intensity of the spot in donor (green) and acceptor (red) channels, including baseline intensities directly after the trajectory where available; Lower: Corresponding FRET efficiency for duration of each trajectory. (C) Image sections of acceptor (upper) and donor (lower) channels at key points in a trajectory shown in (B,vi), each showing 3 consecutive frames: first 3 frames of the movie; acceptor photobleaching event; last 2 frames of the trajectory. Frame time: 20 ms. Image shown is 25 *μ*m x 25 *μ*m. White square indicates spot location. (D) All spots FRET efficiency with gaussian fit (excluding datapoints after acceptor photobleaching event), calculated from 11 trajectories (E) Mean squared displacement vs. observation time with linear fit from SPT of all spots in the acceptor channel. Calculated from 20 trajectories.

### Single-molecule folding of OmpG from urea

Next we examined folding of OmpG-Cy3Cy5 from urea. During image acquisition, OmpG-Cy3Cy5 (8M Urea, 250 mM NaCl, 25 mM Tris-HCl pH 7.0) was added to the solution above a preformed DCPC SLB, yielding a dilution to 0.1 M Urea and a final L:P of 1.5 × 10^6^ (Figure 4A). Using the same trajectory selection criteria as with pre-folded OmpG, we identified trajectories of high-FRET spots from the point of their arrival at the bilayer.

**Figure 4.**
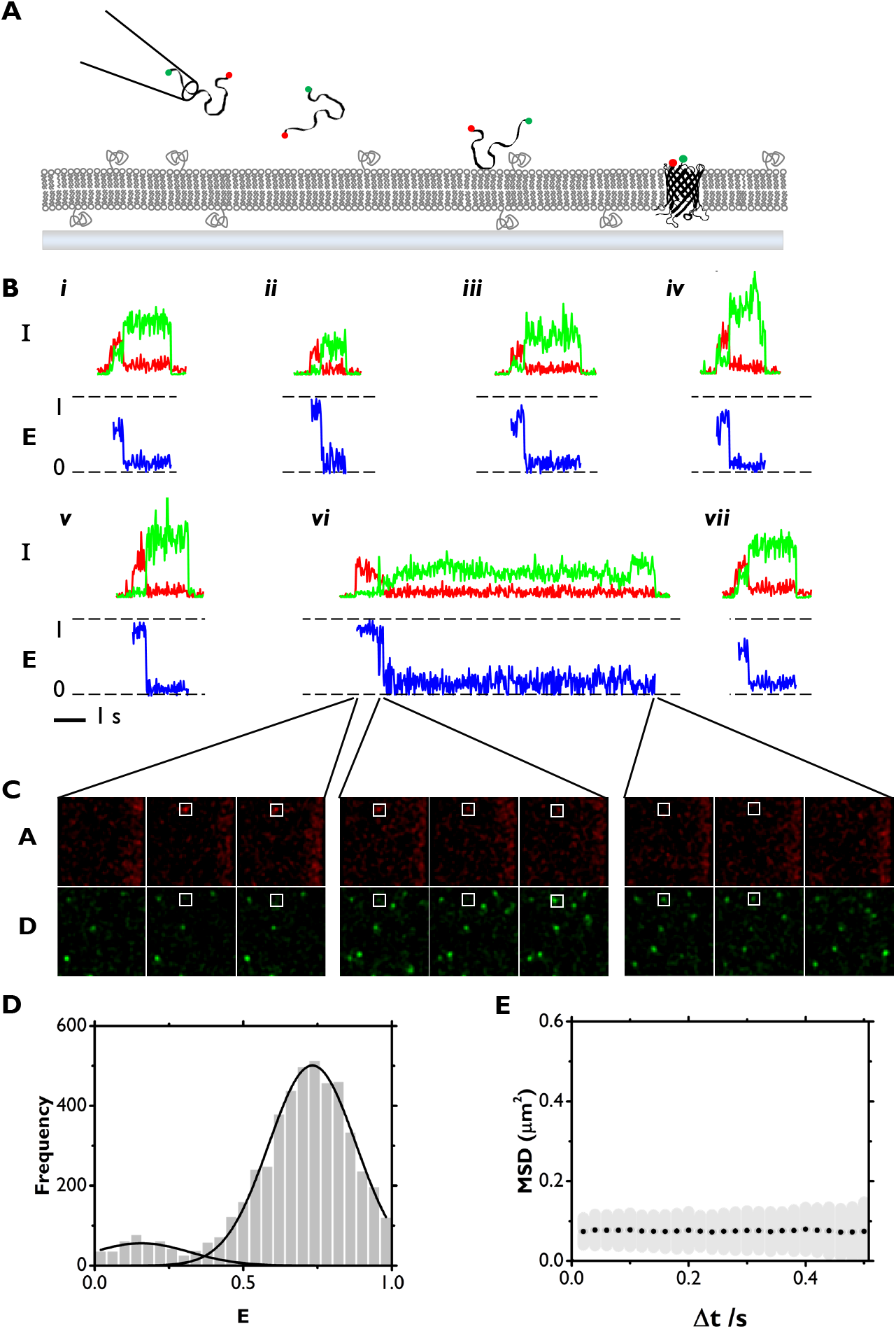
Spontaneous folding and insertion of OmpG in DCPC SLBs: smFRET and SPT. (A) Urea-unfolded OmpG-Cy3Cy5 was injected into the bulk solution surrounding a DCPC SLB. The injection coincided with a rapid dilution of urea, allowing for spontaneous folding and insertion. (B) Representative trajectories of tracked spots. Upper: Intensity of the spot in donor (green) and acceptor (red) channels, including baseline intensities directly before and after the trajectory; Lower: Corresponding FRET efficiency for duration of each trajectory. (C) Image sections of acceptor (upper) and donor (lower) channels at key points in a trajectory shown in (B,vi), each showing 3 consecutive frames: first 2 frames of the trajectory; acceptor photobleaching event; last 2 frames of the trajectory. Frame time: 20 ms. Image shown is 25 *μ*m x 25 *μ*m. White square indicates spot location. (D) All spots FRET efficiency (excluding datapoints after acceptor photobleaching event) with gaussian fit, calculated from 53 trajectories (E) Mean squared displacement vs. observation time with linear fit from SPT of all spots in the acceptor channel. Calculated from 169 trajectories

The majority of trajectories (91%) showed high (E > 0.5) FRET efficiencies for their entire duration (Figure 4B). A wider range of FRET efficiencies than for prefolded OmpG was also observed; evident from the shifting (〈E〉 = 0.73) and broadening of the high-FRET population (Figure 4D). A small subset of trajectories (9%) contained events corresponding to fluctuations at low (0-0.5) FRET efficiency values (Figure 5D) not observed with pre-folded OmpG (Figure 4D). These rare, fluctuating events are a likely cause for the differences observed in average FRET efficiency between pre-folded and injected OmpG-Cy3Cy5. After 10 minutes, fluctuations between high and low FRET states were no longer observed and arrival of new OmpG molecules to the bilayer had ceased (Figure S4).

**Figure 5.**
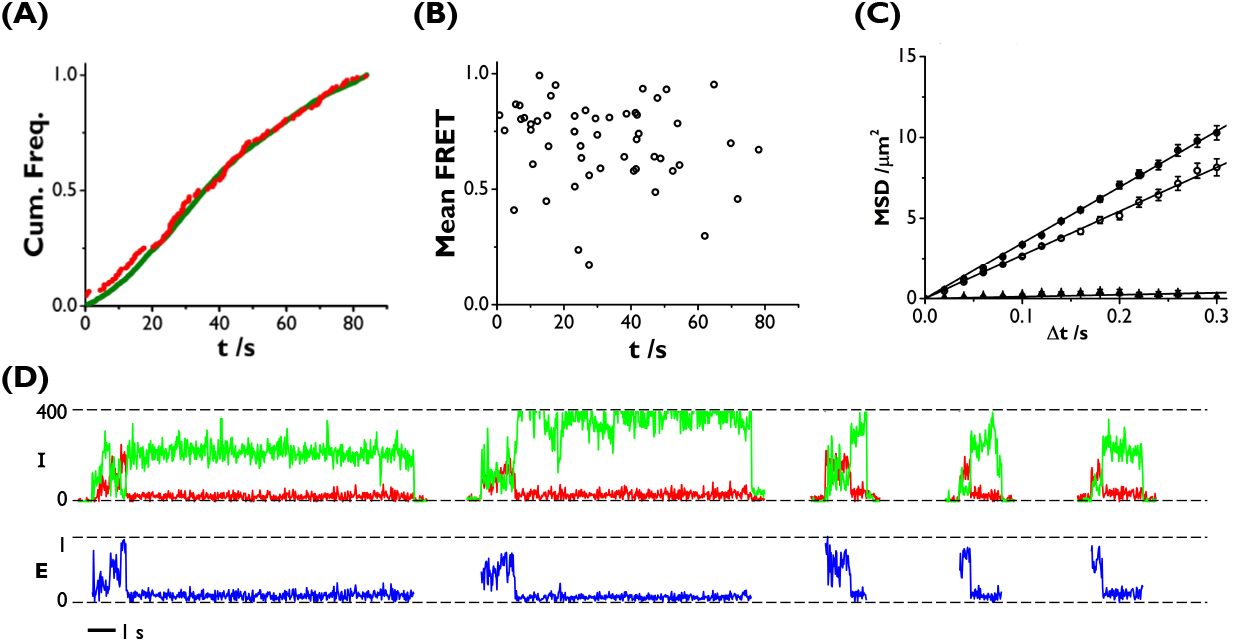
Arrival rates and rare events. (A) Spot arrival times of injected OmpG-Cy3Cy5 landing on a DCPC SLB from a urea-unfolded state, detailed in figure 3. Green: spots detected in the donor channel (3596 in total). Red: Spots detected in the acceptor channel (169 in total) (B) Mean FRET vs. start time for high-FRET trajectories of injected OmpG-Cy3Cy5 in DCPC SLBs. (C) Mean squared displacement vs. observation time interval with linear fit for SPT of fluorophore-labelled lipids in DCPC SLBs before (filled circles) and 10 minutes after (empty circles) addition of urea-unfolded OmpG Cy5 (diamonds) (D) Trajectories of injected OmpG-Cy3Cy5 which have unusual (not consistently greater than 0.5) FRET efficiency encountering the DCPC bilayer.

In contrast to the pre-folded OmpG-Cy3Cy5, following renaturation from urea OmpG-Cy3Cy5 were essentially immobile at the bilayer (Figure 4E). The rate at which OmpG-Cy3Cy5 arrived at the bilayer in our measurements was constant for approximately 50 seconds after injection and a majority of spots were found to have arrived during the 84 s measurement (Figure S5) reflecting the time taken for a majority of OmpG molecules to diffuse from the pipette tip to the SLB; a distance of 2-3 mm. Although the frequency of low-FRET OmpG-Cy3Cy5 arriving at the bilayer was 20-fold greater than the high-FRET trajectories, the cumulative frequency distributions of both are identical, indicating that arrival rate is the same regardless of observed folded state (Figure 5A).

## Discussion

Overall, our experimental evidence supports a simple two-state folding model for OmpG, absent from folding intermediates. Our band-shift assays and ensemble FRET yield folding rates identical within experimental error. smFRET reveals just a single folded state, consistent with what Rath and co-workers similarly observed for OmpX [78] and PagP [79].

Consistent with previous reports, our OmpG ensemble measurements indicated folding kinetics on the order of several tens of seconds. These kinetics were reflected in single molecule measurements where the arrival rate of OmpG at the bilayer occurred on a similar time scale, with a majority of the protein being found to arrive during the course of the 84s recording (Fig S5). However, individual folding events observed at the single-molecule level, regardless of arrival time, are orders of magnitude faster (j20 ms).

Single-molecule measurements are limited by photobleaching time; in our case fluorophores bleached 1-10 seconds after arrival at the bilayer. We can therefore only comment on the folding kinetics of proteins which fold within this observation window. 90% of the OmpG that we observed in a folded state were detected immediately in their final folded state. Thus these events must have occurred faster than the time resolution of our measurements; within 20 ms of arrival of an OmpG molecule at the bilayer. Clearly future insights stand to be gained by single-molecule techniques capable of improved temporal resolution.

Using timecourse band-shift assays, Burgess et al. reported folding kinetics for a range of OMPs into DCPC membranes that were on the order of several tens to hundreds of seconds, but they noted that some OMPs displayed a significant folded fraction existed by their first measured timepoint (5 s) [32]. They termed these rapid folding events a ’burst’ phase, which was found to account for up to around 80% of folding for some OMPs (e.g. OmpX) but was not detectable in others. We did not require the inclusion of a burst phase in order to fit OmpG band-shift timecourse data, so conclude that the proportion of OmpG ’burst’ folding must be low. On the other hand, single molecule assays are sensitive to rare events, so we attribute the small number of smFRET folding events that we observed to the rapid folding events described by a ’burst’ phase. Given the infrequent nature of the few events that displayed a fluctuating FRET efficiency (Figure 5D) it is difficult to speculate further, perhaps these represent instances where the protein transitioned to a misfolded state in which the N- and C-termini were in close proximity. Whether or not this transition is reversible is not known. We would not expect to see these rare, short-lived events in the band-shift or bulk FRET measurements, as they would be obscured by averaging of the signal. It is not possible to draw firm conclusions about the nature of a mis-folded state from so few examples, other than to acknowledge the possibility of its existence.

Assuming each OmpG molecule associates with the SLB long enough to be detected once and is distributed evenly throughout the SLB, the expected spot density (0.33 μ*m*^−2^) also closely matches the observed total spot density (0.27 μ*m*^−2^). Comparing total spot density to the spot density in the acceptor channel over the same period, we can estimate the fraction of molecules that fold on arrival at the bilayer, and taking into account the population of FRET-irresponsive donor-only labelled species and assuming that acceptor-only labelled species are not visible, we estimate the probability of insertion of an individual molecule upon encountering the bilayer to be low, ≈ 0.07.

It is known that the rate of folding of OMPs into membranes is influenced by the accessibility of the hydrophobic bilayer interior; more rapid folding is observed for thinner bilayers, those with increased curvature, or those at their transition temperature [32, 80]. We therefore interpret the folding events that we observed are those in which OmpG landed in an orientation and region of the membrane, leading to a rapid (millisecond) folding event. The majority of encounters do not result in a folding event.

In contrast to the pre-folded OmpG-Cy3Cy5, injected OmpG-Cy3Cy5 did not diffuse (Figure 5E), which did not change over the subsequent 10 minutes (Figure S4E). We were concerned that bilayer defects might be the cause of this change in mobility, and thus affect our kinetics: The mobility of the lipids in the SLB was assessed by incorporating a small fraction of fluorophore-labelled lipid and SPT was used to track the 2D diffusion before and after injection of OmpG Cy5 (Figure 5C). The lipids remained mobile, with no evidence of anomalous subdiffusion, where previously we have exploited PEG-induced bilayer defects to control anomalous subdiffusion in SLBs [81]. Here we saw no dependence of lipid diffusion on observation time (Figure S6), indicating that the PEG-cushioned SLBs in this work were, as expected, free from such defects. A second possible cause for this reduced mobility of OmpG-Cy3Cy5 could be interactions between the protein and underlying glass substrate. Previous reports suggest that spontaneous insertion occurs such that N- and C-termini do not traverse the bilayer [24]. We therefore expect that injected OmpG would be orientated with the large periplasmic loops situated between the lower leaflet and glass substrate and the N- and C-termini situated in the bulk solution. In contrast, we speculate that the orientation of pre-folded OmpG-Cy3Cy5, which is dictated by the SUV rupture mechanism, could be the opposite way around, resulting in a substantially smaller proportion of the protein residing below the lower leaflet, and thus higher mobility.

These experiments highlight the utility of single-molecule tools for dissecting difficult biological processes such as membrane protein folding. Fast, infrequent, folding provides a consistent simple explanation bridging both ensemble and single-molecule observations of kinetics. Here we have deliberately chosen OmpG as a simple testbed with ’uncomplicated’ kinetics and structure; the real challenge is to bring these tools to bear to a wider array of complex challenges in membrane protein folding.

