## Supplementary Information for "Fast slow folding of an Outer Membrane Porin"

### Fast slow folding of an Outer Membrane Porin. Supplementary Information.

Compiled 6 April 2021

**Abstract.** In comparison to globular proteins, the spontaneous folding and insertion of $\beta$-barrel membrane proteins is surprisingly slow, typically occurring on the order of minutes. Using single-molecule Förster Resonance Energy Transfer to report on the folding of fluorescently-labelled Outer Membrane Protein G we measured the real-time insertion of a $\beta$-barrel membrane protein from an unfolded state. Folding events were rare, and fast (<20 ms); occurring immediately upon arrival at the membrane. This combination of infrequent, but rare, folding resolves this apparent dichotomy between slow ensemble kinetics, and the typical timescales of biomolecular folding.

#### Extended Materials and Methods

*Materials*

1,2-dicapryl-*sn*-glycero-3-phosphocholine (DCPC) and Diphytanoylphosphatidylcholine (DPhPC) were purchased from Avanti Polar Lipids (Alabaster, AL). Cy3-maleimide and Cy5-maleimide were purchased from Fisher Scientific. 1,2-dipalmitoyl-*sn*-glycero-3-phosphoethanolamine-N-[methoxy(polyethylene glycol) - 5000] ammonium salt (PEG(5K)-DPPE) and Texas Red 1,2-dihexadecanoyl-*sn*-glycero-3-phosphoethanolamine triethylammonium salt (TR-DHPE) was purchased from Lipoid (Ludwigshafen, Germany). n-Dodecyl-$\beta$-D-Maltopyranoside (DDM) was purchased from Anatrace. Unless stated, all other chemicals were purchased from Sigma-Aldrich. All aqueous solutions were prepared using doubly deionized 18.2 MΩcm MilliQ water.

*Expression, purification and labelling of OmpG-2xCys*

The plasmid pT7-OmpGwt was used as the template [**?**] for a double cysteine mutant of full-length OmpG, produced by introducing two cysteine residues at positions E2C and C281 using a multimutagenesis kit (Stratagene). See Table S1 for primer sequences.

Labelling sites at residues 2 and 281, near the N- and C-termini, were selected in order to optimise the change in FRET efficiency on transitioning between unfolded and folded states. In a folded state, the $C_\alpha$ atoms of the labelling sites are 1.1 nm apart according to the crystal structure of detergent-folded OmpG. In an unfolded state, the residues are free to diffuse independently in solution, tethered by a 279-residue polypeptide chain, with a predicted mean separation of 15.2 nm. The Cy3-Cy5 FRET pair used here has a Förster radius sensitive to this distance range ($R_0$=5.4 nm). Residues in the $\beta$-barrel structure or on extracellular loops, which are likely to penetrate the bilayer, were avoided.

Constructs were expressed and purified using a previously described method [**?**]. Briefly, pT7-OmpG (E2C X281C) plasmid was transformed into *E. coli* PC2889 cells [BL21(DE3)Δ lamB ompR] [**?**] and protein expression was induced by the addition of IPTG under the control of the T7 promoter. Cells were harvested and OmpG-2xCys was extracted from inclusion bodies using a denaturing buffer [8M Urea, 1 mM TCEP, 50 mM Tris.HCl, pH 8.0 ] and purified by anion exchange. The purified OmpG-2xCys was either labelled or stored at -80$^o$C until required. OmpG-2xCys was then purified by gel filtration [eluting in 8M Urea, 250 mM NaCl, 25 mM Tris.HCl (pH 8.0)] in order to remove TCEP. Labelling was performed immediately afterwards by incubation with a 10-molar excess each of Cy3- and Cy5-maleimide for 60 min at room temperature. Excess label was removed by gel filtration.

*Labelling efficiency*

The average labelling efficiency of OmpG Cy3/Cy5 was determined by separately quantifying OmpG and both fluorophores by absorption spectroscopy using a NanoDrop

8000 Spectrophotometer (Thermofisher Scientific). See Table S2 for all of the above calculations and figure S1 for spectra. Concentrations were calculated using the Beer-Lambert law, $A = \varepsilon c l$, where $A$ is the absorption due to that particular species, $\varepsilon$ is the absorption coefficient (see Table S1) and $l$ is the pathlength. For Cy5, the absorption at the wavelength where it is most strongly absorbing ($\lambda_{\mathrm{max}}$) is used without any correction:

$$A_{\mathrm{Cy5}} = A_{652\mathrm{nm}}. \tag{1}$$

For Cy3, correction is required to account for the contribution from Cy5 at $\lambda_{\mathrm{max}}$,

$$A_{\mathrm{Cy3}} = A_{550\mathrm{nm}} - A_{\mathrm{Cy5}} \times \mathrm{CF}_{\mathrm{Cy5}}, \tag{2}$$

where the correction factor ($\mathrm{CF}_{Cy5}$) was determined from the absorption spectrum of free Cy5 in solution as $A_{550\mathrm{nm}}/A_{652\mathrm{nm}}$. For OmpG, correction is required to account for the contribution from both fluorophores at 280 nm,

$$A_{\mathrm{OmpG}} = A_{280\mathrm{nm}} - (A_{\mathrm{Cy3}} \times \mathrm{CF}_{\mathrm{Cy3}}) - (A_{\mathrm{Cy5}} \times \mathrm{CF}_{\mathrm{Cy5}}), \tag{3}$$

where the correction factors were determined from the absorption spectra of free fluorophores as $A_{280\mathrm{nm}}/A_{\lambda\mathrm{max}}$. The labelling efficiency ($E$) is calculated as:

$$E = \frac{[\mathrm{Dye}]}{[\mathrm{OmpG}]}. \tag{4}$$

The labelling efficinecy of OmpG-Cy3Cy5 was 0.98 and 1.00 for Cy3 and Cy5.

*Electrophysiology*

For single-channel recordings, OmpG-Cy3Cy5 was refolded in detergent micelles and then reconstituted in a droplet interface bilayer (DIB). For detergent refolding, 5µl OmpG-Cy3Cy5 stock was mixed with 245 µl refolding buffer (1 M KCl, 10 mM Tris pH 7.2, 0.5% Genapol X-080) and incubated at 37$^{o}$C for 2 hours. Refolded OmpG-Cy3Cy5 was diluted to 100 pM in KCl Buffer (1M KCl, 10 mM Tris pH 7.2) immediately before use. Droplet interface bilayers were produced as described in [?]. Briefly, 100 µl molten agarose in water (0.75 %, 90oC) was spin-coated (3000 rpm, 30 s, using a Laurell Technologies spin coater) on to an $O_2$ plasma-cleaned glass coverslip, providing a thin (¡ 100 nm) hydrogel layer. The coverslip was assembled with a microfabricated PMMA device comprising a planar fluidic channel contacting the underlying agarose base layer, interdigitating an array of vertical open wells which allow access to the spin-coated agarose from the top of the device. This channel was filled with a molten solution of agarose (2.5% w/v, 90$^{o}$C) in KCl buffer filling the void of the channel, providing a source of hydration to the underlying agarose layer while maintaining it's suitably thin depth for evanescent field penetration in subsequent TIRF imaging. A Ag/AgCl electrode was inserted into the rehydrating agarose while still molten. The agarose layer within each well was submerged in oil solution (hexadecane:silicone oil AR20 (9:1)) containing dissolved DPhPC at 8.5 mg mL$^{-1}$) and allowed to equilibrate for 30 min. A 20 nL aqueous droplet of refolded OmpG-Cy3Cy5 diluted in KCl buffer was incubated in a separate chamber containing the same lipid in oil solution for 20 minutes before being transferred by pipette to the experimental device. A bilayer was formed upon the droplet sinking to the bottom of oil-containing well and making contact with the underlying agarose surface. An agarose-coated Ag/AgCl electrode was inserted into the droplet by making contact from above, using a micromanipulator. Both electrodes were connected to the headstage of an Axopatch 200B patch-clamp amplifier (Molecular Devices) which was interfaced with a computer via a BNC 2090A connector block and an NI PCI e-6251 M-series data acquisition card (both National Instruments). Recordings of ionic current were made in voltage-clamp mode at room temperature and digitized at a rate of 1 kHz. A gaussian filter ($\sigma = 5$, $\mu = 30$) was applied to the measured current.

*SUV preparation*

To prepare SUVs, DCPC dissolved in chloroform was dried under a stream of $N_2$ gas and then under vacuum for 1 hour. The dried lipids were hydrated with water or buffer and vortexed before tip sonication (Vibracell VCX130PB with CV188 tip, Sonics & Materials, Newtown, CA) for 15 minutes at 25% amplitude. The resulting clear vesicle suspension was centrifuged (3 minutes; 14000 × $g$) before the supernatant was retained and any titanium residue (from the sonicator probe) was discarded. SUV preparations were stored at 4°C for up to 48 hours. Where required, PEG(5K)DPPE and TR-DHPE were incorporated by dissolving in chloroform and then mixing with DCPC (in chloroform) prior to drying down.

*Bulk folding assay*

For bulk folding assays, DCPC SUVs were prepared at 2.5 mg/ml in NaCl Buffer (250 mM NaCl, 10 mM Tris pH 7, 10 mM DTT). OmpG-2xCys or OmpG-Cy3Cy5 was mixed with SUVs and NaCl Buffer to a total volume of 15 µl, diluting the urea in the protein sample from 8M to 0.8 M. The ratio of denaturant to renaturant (e.g. lipid vesicles) controls the equilibrium between folded and unfolded states [**?**]. Samples were incubated at 37°C before quenching with 5µl SDS Buffer (250 mM NaCl, 10 mM Tris pH 7, 10 mM DTT, 0.8 M Urea, 1% (w,v) SDS). For the timecourse assay, L:P was fixed at 6250 and incubation time was varied from 0 - 60 min. For the titration, all samples were incubated for 60 min, and L:P was varied from 0 - 6803. Emission spectra were obtained at room temperature with excitation separately at 532 nm and 600 nm using a Cary Eclipse Fluorescence Spectrophotometer (Agilent). For the band-shift assay, 15 µl of each sample was mixed with 5 µl RunBlue 4x LDS Sample Buffer (Expedeon) and 15 µl was loaded onto a 10 % Run-Blue SDS polyacrylamide gel (Expedeon) for electrophoresis. Gels were imaged using an Amersham Imager 600 (GE Healthcare). For FRET SDS-PAGE, gels were excited with 520 nm light and imaged separately through 605/40 BP (donor) or 705/40 BP (acceptor) emission filters. For densitometry analysis, gels were excited with 630 nm light and imaged through a 705/40 BP emission filter in order to obtain a FRET-independant signal by directly exciting the acceptor fluorophore only. For OmpG-2xCys, gels were stained with InstantBlue (Expedeon) and then imaged with white light.

*Bulk fluorimetry*

To calculate FRET efficiency, emission spectra were first normalised for variations in sample concentration (determined by maximum emission at 600 nm excitation) and then FRET-independent signals were subtracted. The magnitude of the FRET-independent contributions was derived from an emission spectrum of OmpG-Cy3Cy5 after proteolysis by proteinase K, and knowledge of the OmpG-Cy3Cy5 labelling efficiency (Figure S3). In the 'zero FRET' state, it is assumed that all polypeptide chains connecting fluorophores are proteolytically cleaved, therefore any interaction between donor and acceptor fluorophores is at random and the FRET efficiency is 0. The FRET-independent signal from donor-only labelled OmpG species was derived from the spectrum of free Cy3 in solution normalised to 0.5 × the donor peak height in the proteinase K digested OmpG-Cy3Cy5 sample. The two FRET-independent donor and acceptor signals were subtracted from each concentration-normalised spectrum, and the relative FRET efficiency ($E$) was calculated as

$$E = \frac{A}{A + D} \tag{5}$$

where $A$ and $D$ are the emission values at 567 nm and 667 nm.

*SDS-PAGE Image analysis*

Fluorescent SDS PAGE images were processed using ImageJ (Figure S2). Donor ($D_{\text{gel}}$) and acceptor ($A_{\text{gel}}$) images were cropped to display identical regions and contrast was adjusted to display the full range of band intensities before applying a black-green (donor) or black-red (acceptor) linear lookup table. The FRET gel image ($E_{\text{gel}}$) was produced from donor and acceptor images

$$E_{\text{gel}} = \frac{A_{\text{gel}}}{A_{\text{gel}} + D_{\text{gel}}}. \tag{6}$$

A yellow-black linear lookup table was applied, to pixels over the range 0.2 to 0.3.

For band-shift assays, densitometry analysis was carried out using the Gel Analyzer plugin for Image J. The folded fraction ($\alpha$) is calculated as

$$\alpha = \frac{F}{F + U} \tag{7}$$

where $F$ and $U$ is the area under the curves corresponding to folded and unfolded OmpG bands.

*Supported lipid bilayers*

SLBs were prepared on glass coverslips by vesicle fusion [?] from SUVs consisting of 1.77 mM DCPC with 1.0 mol% PEG(5K)-DPPE (and 3x$10^{-6}$ mol% TR-DHPE for lipid tracking experiments). Glass coverslips were rigorously cleaned using stepwise bath sonication with DECON-90, water, and propan-2-ol for 20 minutes each. Immediately before use, the glass was dried under nitrogen and cleaned with oxygen-plasma treatment for 3 minutes (Diener Electronic, Femto). A 1 cm (approx.) diameter well was created on each coverslip using vacuum grease (Dow Corning). The coverslip was heated to 37°C before 50 µL of SUV stock were diluted 1:1 in SLB Buffer (250 mM NaCl, 10 mM EDTA, 10 mM Tris pH 7.0) and added to the chamber immediately. The vesicles were incubated for 30 minutes before the membranes were washed thoroughly with water followed by NaCl buffer (250 mM NaCl, 10 mM Tris pH 7.0) maintaining the volume at 100-150 µl. Samples were heated to 37$^o$C throughout imaging, using a custom-made objective heater. For injection measurements, OmpG-Cy3Cy5 was diluted to 7 pM in denaturing buffer (8M urea, 250 mM NaCl, 10 mM tris pH 7) then 2 µl was added to the bulk solution during image acquisition. For SLBs containing pre-folded OmpG-Cy3Cy5, the protein stock was diluted to 700 pM in denaturing buffer, then 5 µl was mixed with 45 µl SUV stock and incubated for 1 hour at 37$^o$C prior to SLB formation. For lipid tracking measurements, SLBs were imaged before and 10 minutes after the addition of OmpG Cy5, as described for OmpG-Cy3Cy5 injection.

*Microscopy*

A 532 nm continuous-wave laser was focussed at the back aperture of an objective lens (60× TIRF oil-immersion NA 1.49, Nikon, ∼0.9 kW $cm^{-2}$) mounted on an inverted microscope (Ti Eclipse, Nikon Instruments, UK) giving rise to total internal reflection at the coverslip surface. Emitted fluorescence was collected through the same objective, transmitted through a red/green dual-dichroic mirror (XF2055, Horiba) prior to passing through a long pass emission filter (550 nm, Chroma) followed by wavelength image splitting to separate out donor and acceptor emission for side-by-side imaging on a single 128 × 128 pixel frame-transfer emCCD detector (iXon DU-860, Andon Technology PLC, Belfast, UK). Wavelength image splitting was achieved with an Optosplit II (Cairn Research, UK) using a dichroic beamsplitter (FF660 Semrock, USA) and bandpass filters in the donor (550/88 Semrock) and acceptor (680/42 (Semrock), 664 LP (Edgebasic)) channel light paths, selected to minimise donor and acceptor emission bleed. For lipid tracking measurements, the labelled lipids were imaged using the donor channel of the above setup. Images were acquired with a frame time of 20 ms and EM gain of 300 and saved as 16-bit Tagged Image File Format (TIFF) for subsequent analysis. A beam shutter TTL synchronised to the emCCD ensured that the imaged region experienced no laser exposure prior to image acquisition to avoid photobleaching. Data were a acquired from a minimum of 3 SLBs per experiment. For pre-folded, post-injection and labelled lipid experiments, multiple time series acquisitions of 30 s were recorded for each bilayer, with each acquisition in a previously unexposed area of the bilayer. For injection experiments, a maximum of 2 time series acquisitions of 100 s were recorded per bilayer, with the injection occurring within the first 5 s. For the second injection recording, a new area of the bilayer was exposed to the laser in order to photobleach all fluorophores in the field of view prior to the acquisition and subsequent injection.

*Image analysis*

All data were analysed using Matlab (R2017b, MathWorks USA) unless stated. To prepare image stacks for spot detection and tracking, a gaussian blur was applied to all frames ($\sigma = 10$) and background subtraction was performed via the pixel-wise subtraction of time averaged median pixel intensity from each image sequence. Image stacks were subsequently split into donor and acceptor channel images ($64 \times 128$ pixels). The affine transformation required to register acceptor to donor channel was derived from graticule images, taken under the same optical configuration as the data during data acquisition, using the Descriptor-based Registration plugin for ImageJ [**?**]. The transformation was applied to the acceptor image, using cubic interpolation for smooth rendering. For diffusion analysis, SPT was performed using TrackMate [**?**], a plugin for ImageJ. For lipid tracking diffusion analysis, the space-time co-ordinates of the output tracks were used to calculate mean-squared displacements for different observation times using a custom-written procedure in Matlab (MathWorks) as described previously [**?**].

Trajectories of folded OmpG-Cy3Cy5 were initially obtained using TrackMate. A plot of maximum donor and acceptor intensity was produced, for the duration of the entire recording, within a square region of interest (ROI) about initial trajectory location. Start and end frames were assigned manually, based on visibility of spots in the donor channel. Trajectories were retained if three conditions were satisfied: anticorrelation of donor and acceptor channels; a single photobleaching step in each channel; an absence of neighbouring spots in the ROI during the trajectory. Precise spot locations were obtainied by a 2D gaussian fitting routine on the summed donor and acceptor images. Spot intensities were given as maximum pixel intensities from each channel within a 3 x 3 pixel region about the spot location. FRET efficiencies for each spot ($E_{\text{spot}}$) were extracted frame-by-frame from the donor ($D_{\text{spot}}$) and acceptor ($A_{\text{spot}}$) spot intensities as follows:

$$E_{\text{spot}} = \frac{A_{\text{spot}}}{A_{\text{spot}} + D_{\text{spot}}}. \tag{8}$$

#### Supplementary Tables

| Mutation | Sequence |
|---|---|
| E2C | *Forward:TATACATATGTGCGAAAGGAACGACTGG* |
| | *Reverse:TTCCTTTCGCACATATGTATATCTCCTTC* |
| C281 | *Forward:TACTCGTTCTGTTGATAAAAGCTTGGATC* |
| | *Reverse:TTTTATCAACAGAACGAGTAATTTACGCC* |

**Table 1. Table S1: Mutagenesis primer sequences**

| Species | $\lambda_{\mathbf{max}}$ (nm) | CF | $\varepsilon$ (μM$^{-1}$cm$^{-1}$) | $A_{\lambda\max}$ | $A_{\text{species}}$ | C (μM) | E |
|---|---|---|---|---|---|---|---|
| Cy5 | 652 | $0.03^a$ $0.05^b$ | $250,000^c$ | 0.177 | 0.177 | 7.08 | 0.98 |
| Cy3 | 550 | $0.04^a$ | $150,000^c$ | 0.117 | 0.108 | 7.21 | 1.00 |
| OmpG | 280 | N/A | $87,905^d$ | 0.073 | 0.063 | 7.18 | N/A |

**Table 2.** Calculation of OmpG Cy3/Cy5 labelling efficiency. Values taken from the spectra shown in Figure S1, unless stated otherwise. $\lambda_{\max}$: Wavelength of absorption peak. CF: Correction factor denoting contribution of the species to the absorption at (a) 280 nm and (b) 550 nm as a proportion of the absorption at $\lambda_{\max}$. $\varepsilon$: Extinction coefficient obtained (c) from the manufacturer or (d) based on the amino acid sequence, calculated using ExPasy ProtParam [**?**]. $A_{\max}$: Absorption at $\lambda_{\max}$. $A_{\text{species}}$: Absorption at $\lambda_{\max}$ due to that particular species. $C$: Concentration. $E$: Labelling efficiency.

#### Supplementary References

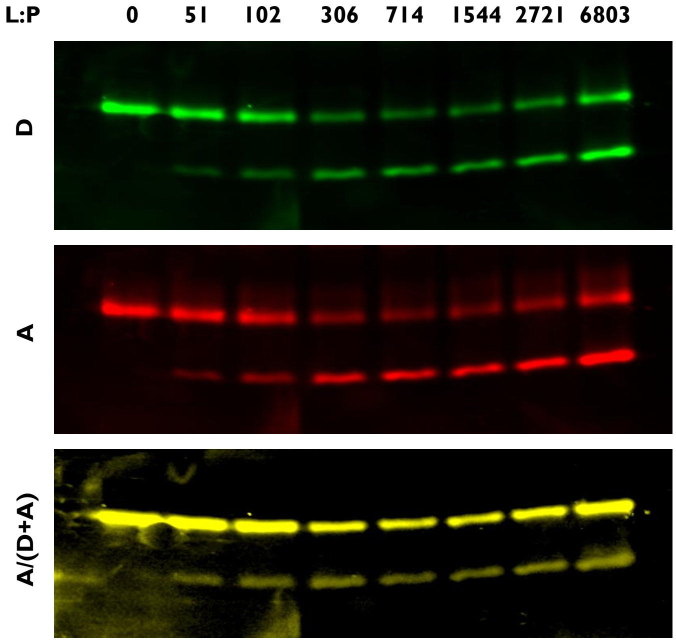


**Figure 1. Image Processing of Fluorescent SDS PAGE** SDS-PAGE of urea-unfolded OmpG-Cy3Cy5 after 1 h incubation with DCPC SUVs in 0.8 M urea, at varying L:P. The gel was illuminated at the donor fluorophore absorption wavelength, and imaged separately through emission filters corresponding to donor ('D' - top) and acceptor ('A' - middle) emission wavelengths. The bottom image is the acceptor emission divided by the sum of the donor and acceptor images, scaled linearly from 0.2 (yellow) to 0.3 (black).

#### Supplementary Figures

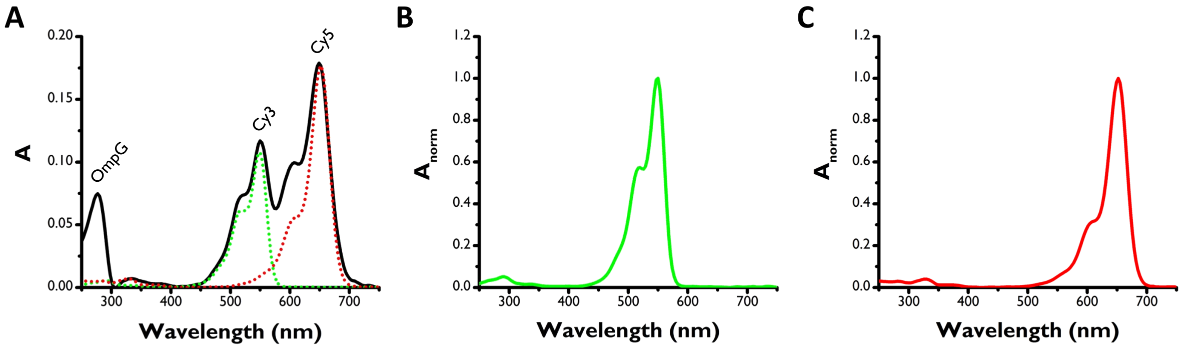


**Figure 2. OmpG Cy3/Cy5 Absorption Spectrum** (A) OmpG-Cy3Cy5 (Black) with normalised Cy3 maleimide (green) and Cy5 maleimide (red) to show contributions from individual species. Free Cy3 maleimide (B) and Cy5 maleimide (C). All samples in OmpG purification buffer.

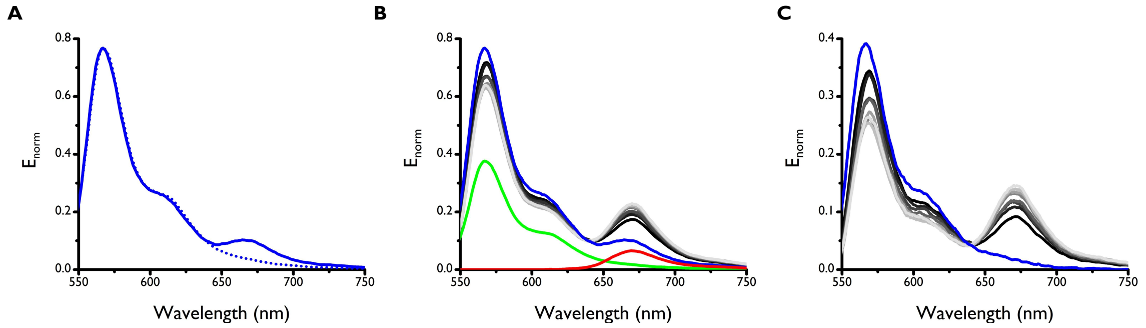


**Figure 3. Processing OmpG-Cy3Cy5 emission spectra** Emission spectra of OmpG-Cy3Cy5 folding into DCPC SUVs timecourse from 0 (black) to 60 (grey) minutes and OmpG-Cy3Cy5 after digestion by protease K (blue): (A) with normalised Cy3 emission spectrum (dashed line); (B) with donor (green) and acceptor (red) FRET-independent signals; and (C) after subtraction of FRET-independent signals. Sample excited at 532 nm. Spectra normalised for variations in concentration (see methods section). The FRET-independent acceptor emission, presumably caused by direct excitation of the fluorophore (whose absorbance is approx. 4 % of its maximum at 532 nm) is the difference between the digested OmpG-Cy3Cy5 spectrum and the normalised Cy3 spectrum. The FRET-independent donor emission was derived by multiplying the Cy3 spectrum by the expected population of donor fluorophores in solution that do not share an OmpG polypeptide with an acceptor fluorophore. Given the labelling efficiency of OmpG-Cy3Cy5 and assuming the fluorophores bind both labelling positions with equal preference, on average, 0.5 of the OmpG population is bound by one of each fluorophore, and 0.25 each of either two donor or two acceptor fluorophores. Thus, 0.5 of all donor fluorophores in the sample are not connected to an acceptor fluorophore, and their emission is not expected to change as the proximity of the labelling sites varies

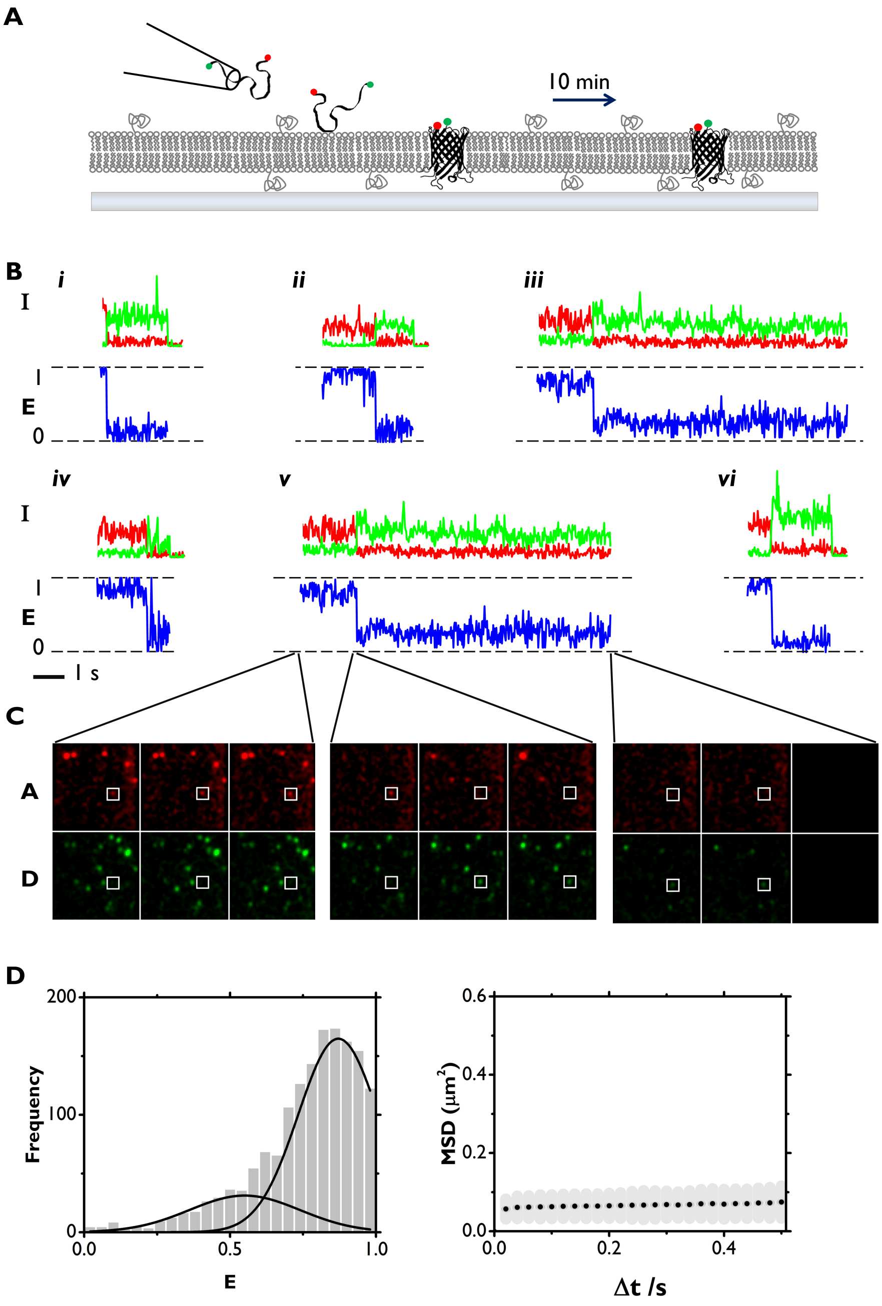


**Figure 4. smFRET of OmpG 10 minutes after spontaneous insertion into DCPC SLB** (A) Urea-unfolded OmpG-Cy3Cy5 was injected into the bulk solution surrounding a DCPC SLB and then incubated at 37°C for 10 minutes. The injection coincides with a rapid dilution of urea, allowing for spontaneous folding and insertion. (B) Representative trajectories of tracked spots. Upper: intensity of the spot in donor (green) and acceptor (red) channels. Lower: corresponding FRET efficiency for the duration of each trajectory. We attribute the broad peak centred at 0.5 to increased background noise, owing to the higher spot density of this dataset in comparison to the previous two. (C) Image sections of acceptor (upper) and donor (lower) channels at key points in a trajectory shown in (B,vi), each showing 3 consecutive frames: first 3 frames of the movie; acceptor photobleaching event; last 2 frames of the trajectory. Frame time: 20 ms. Image shown is 25 $\mu$m x 25 $\mu$m. White square indicates spot location. (D) All spots FRET efficiency (excluding datapoints after acceptor photobleaching event) with gaussian fit, calculated from 43 trajectories. (E) Mean squared displacement vs. observation time with linear fit from tracking spots in acceptor channel, calculated from 167 trajectories.

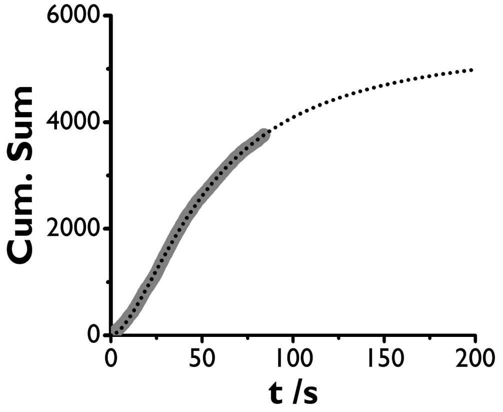


**Figure 5. Arrival rate of injected OmpG-Cy3Cy5 landing on a DCPC SLB from a urea-unfolded state.** Cumulative sum of total spots detected in both donor and acceptor channels over time (3765 over 84 s) with sigmoidal fit extrapolated to 200 s. Plateau = 5547, t = 53.7 s at the midpoint.

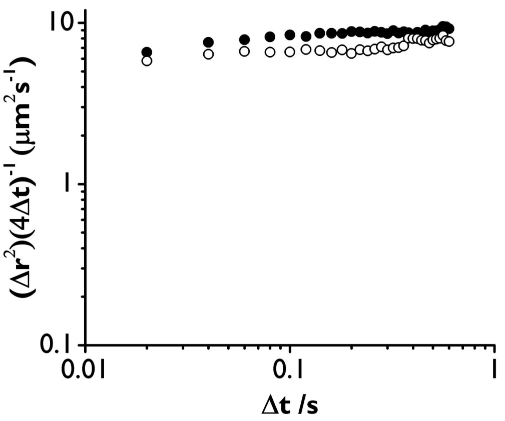


**Figure 6. Anomalous diffusion plot of DCPC SLB** Diffusivity of fluorophore-labelled lipid in DCPC SLB, calculated at varying observation times. Images acquired before (filled circles) and after (open circles) injection of OmpG Cy5.

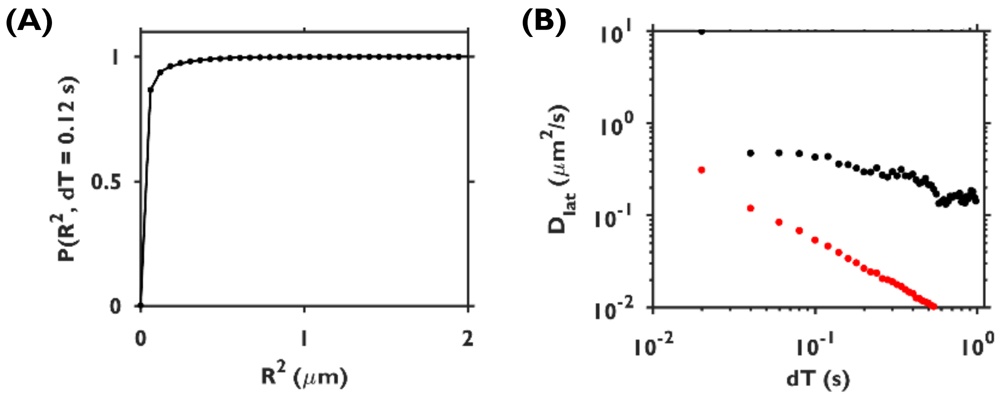


**Figure 7. Two-component diffusivity analysis of low-FRET spots of urea-unfolded OmpG Cy3Cy5 on arrival at DCPC SLB** (A) Probability analysis of radial diffusion for the observation period of 0.12s, with double exponential fit. Dlat = 0.05 µ$ms^{-1}$ for 88% of the population, and 0.43 µ$ms^{-1}$ for 12% at this observation period. (B) Diffusivity at varying observation times for the larger (black) and smaller (red) populations.
